# Study on the interaction of Yohimbine with duplex oligonucleotide using spectroscopic and computational tools

**DOI:** 10.1101/2025.07.31.667832

**Authors:** Soching Luikham, Vibeizonuo Rupreo, Senchumbeni Yanthan, Jhimli Bhattacharyya

## Abstract

DNA-interactions with multivalent ligand(s) have increasingly become the subject of substantial research. For several small-molecules with therapeutic-potential, nucleic-acids serve as their primary molecular-target. Such interaction has been shown to affect transcription, and replication, ultimately leading to apoptotic cell-death. Thus, researchers are becoming increasingly interested in understanding ligand-interaction with oligonucleotides making it possible to develop new, DNA-specific drugs. Yohimbe (Yh), a bioactive indole-alkaloid, has been thoroughly investigated for its pharmaceutical qualities, but the mechanism of DNA-binding is still ambiguous. This research adopted computational and multi-spectroscopic methods to investigate the molecular-mechanism between Yohimbine and hairpin-duplex oligonucleotide at physiological-conditions. The occurrence of slight hypochromic and bathochromic deviations in fluorescence intensity indicates that Yh interacts with hairpin-duplex. Employing the McGhee-von Hipple approach, the Scatchard-plot analyses indicated non-cooperative interaction with 10^5^ M^-1^ binding affinities. The temperature-dependent fluorescence data suggested positive entropy and negative enthalpy supporting the exothermic binding. Salt-dependent fluorescence revealed that non-polyelectrolytic forces governed the DNA-ligand association. The findings of the urea-denaturation, dye-displacement, and molecular-docking analysis, iodide-quenching confirmed groove-binding. Therefore, biophysical tools and *in silico* modeling were utilized to identify the structural-alteration and energetic-profiling of Yh’s interaction with oligonucleotides which can be employed for the development of DNA-targeted therapeutics.

## Introduction

The largest and most intriguing class of naturally occurring bioactive compounds derived from tryptophan is known as the indole alkaloid family. They can be found in a variety of terrestrial and aquatic sources, [1] and are well-known for their broad range of pharmacological and biological effects, including analgesic, anti-HIV, anticonvulsant, anti-fungal, antibacterial, antioxidant, and plant growth regulator etc [2]. Over the preceding years, interest has been drawn to the combination of Indole alkaloids’ bioactivity and structural diversity [3]. Indole alkaloids have therapeutically significant features, making them important for studying their interactions with biomacromolecules to learn about their molecular transport and metabolism, of such good verse indole alkaloid is Yohimbine (Figure 1) (17α-hydroxyyohimban-16α-carboxylic acid methyl ester; Yh) commonly called as Quebrachine, aphrodine, corynine, and hydroaerogotocin[4–6]are famously observed in the pulp of *Pausinystalia yohimbe* from the Central African tree [7]and is utilized to cure a variety of ailments such as male erectile dysfunction, marijuana abuse, depression, idiopathic orthostatic hypotension, and diabetes mellitus type II [8],[9]. It works by inhibiting pre- and postsynaptic α2-adrenergic receptors, which are primarily effective for addressing erectile dysfunction. Yohimbine can also boost sympathetic outflow from the CNS (central nervous system) and catecholamine release from peripheral sympathetic nerve terminals, in addition to limiting the alpha-adreno receptor. β-carboline has thus made a significant impact with their pharmacological value [10],[11]. Due to their extensive occurrence in nature and lesser toxicity when compared to other synthetically created molecules, these bioactive substances are potential elements in pharmacology and pharmacokinetics.

**Figure 1.**
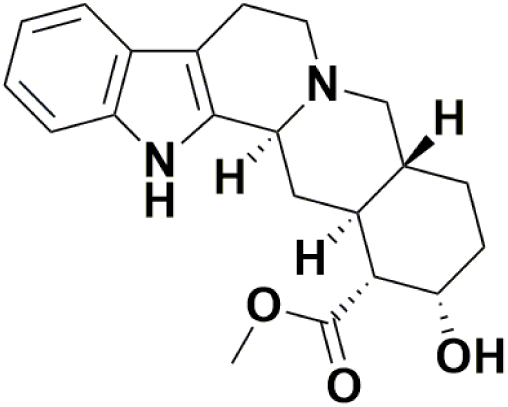
Chemical Structure of Yohimbine (Yh).

Relationships between biological substances and ligands such as deoxyribonucleic acid (DNA), ribonucleic acid (RNA), and protein is a distinguished research topic in establishing biopharmaceutical relevance [12],[13],[14],[15]. DNA plays a significant role in the retrieval of genetic data,[16]which can be a viable treatment option even in cancer and other ailments. Deoxyribonucleic acid serves as the most important bio-macromolecule because it controls the configuration and functioning of the cell. It becomes further an important intrinsic target for anticancer drugs, antibiotics, and other medications [17],[18],[19]. The analysis of DNA as a possible therapeutic threshold is alluring due to the extensively researched 3D structures of DNA and their probable simplicity with numerous essential chemical functional groups. However, in comparison to the presence of protein-based drug targets, DNA-based therapeutic drug research is severely minimal, and the amount of physiologically active DNA-based molecules is supposedly very sparse [20],[21]. Ligand binding to DNA can change the transcription, replication, and expression of genetic information in cells, altering their physiological functioning [22]. Small molecules can interact with DNA through covalent or non-covalent associations, leading to modifications or the inhibition of DNA’s functionality [23]. There are two types of non-covalent contact mechanisms for DNA: intercalation and groove binding, with the thermodynamic profile, binding affinity, and binding modes implicated [24],[25]. The purpose of these investigations is to analyze and identify the binding interactions and pattern specificity of small molecules with GC and AT hairpin duplex oligonucleotides, which are organized by structural components and dynamic electronic, to establish new and effective unique potent treatments for a variety of serious illnesses. The research also could disclose whether Yh preferentially binds to GC or AT base-rich hairpin duplex oligonucleotide sequences [26],[27]. In this research piece; thermodynamic profiling and spectroscopic investigation of Yh association with GC and AT hairpin duplex oligonucleotide using multi-spectroscopic (viz, UV-Vis spectroscopy, Fluorescence spectroscopy, Circular Dichroism (CD), and computational techniques are presented for the first time. This comprehensive study of Yh’s possible therapeutic properties will help researchers build unique and rational medications in the future.

## Materials and methods

### Materials

Yohimbine (Yh) and GC(5’-GCGCGCGTTTTTCGCGCGC3’) and AT(5’-GTATATACCCCCTATATAC-3’) Hairpin duplex oligonucleotides were acquired from the Sigma-Aldrich Corporation. The samples and reactions were prepared using a 10mM sodium cacodylate [NaO_2_As(CH_3_)_2_•3H_2_O] buffer, and they were carried out at a pH of 7.0 which is the physiological pH. Readings of pH and required adjustments were incorporated using a pH meter (digital high-precision) from Systronics. Using molar absorption coefficient (ε) values of 160,100 M^-1^ cm^-1^ at 260 nm for GC and 186,000 M^-1^ cm^-1^ at 260 nm for AT Hairpin duplex oligonucleotide, the concentrations of the analyzed sample were calculated. A standardized stock sample of 1.0 × 10^-3^ Mol L^-1^ of Yh was prepared by dissolving Yh in Millipore water. The necessary chemicals and reagents utilized throughout the present research were of the highest-grade analytical quality from Sigma-Aldrich. Making use of Millipore water, the experimental solutions were made.

### Methods

UV-visible absorption studies were performed at (298.15 K) through an Agilent Cary 100 range UV-Vis spectrophotometer. Fluorescence spectrum analysis, salt-dependent, temperature-dependent, competitive dye displacement, urea denaturation assay, and potassium iodide (KI) quenching were all performed using an Agilent Cary eclipse spectrofluorophotometer. Fluorimetric titration measurements were used to generate data on binding affinity for the Scatchard plot. To measure the CD emission, a JASCO 815 CD spectrophotometer was used. The LCA (Lamarckian Genetic Algorithm) was incorporated in AutoDock 1.5.6 to perform in silico (molecular docking) analysis. The DNA structure was obtained in crystal form the Protein Data Bank with PDB-IDs 1D16 for GC and 5M68 AT for hairpin duplex oligonucleotides. PyMOL (The PyMOL Molecular Graphics System, Version 2.3.4, Schrödinger, LLC) and the UCSF Chimera 1.15 molecular graphics program were used to illustrate each docked orientation. In Supplementary Information (S1) the methods are discussed in detail.

## Results and discussion

### UV-Vis Absorption Spectroscopy

Since Yh has characteristic visible absorption spectra in the 200–300 nm regions with two distinct peaks at 220 nm and 272 nm, respectively, it is feasible to observe the interaction [28],[29]. The electronic shift that occurs within the chromophore present in the pyrimidines and purines was considered to be responsible for both the hairpin duplex oligonucleotide at 258 nm[30]. It was challenging to predict the interaction between Yh and both the GC and AT hairpin duplex oligonucleotide because of the spectral overlap as seen in Figure 2. Upon interacting with GC and AT hairpin duplex oligonucleotide, it was shown that Yh’s UV absorbance at 220 nm decreased, yet both exhibit a small hypochromic effect and a bathochromic shift of 1 nm. This may be because the hairpin duplex base pairs’ π orbitals and the binding ligand’s π*orbitals interact to lower the π-π* transition energy, and cause an absorption redshift. The hypochromic impact was brought on by the electron-filled coupling π *-orbital, which decreased the frequency of transfer [31]. When groove binding and electrostatic interactions occur, only the hypochromic effect can be noticed, regardless of a slight bathochromic effect.

**Figure 2.**
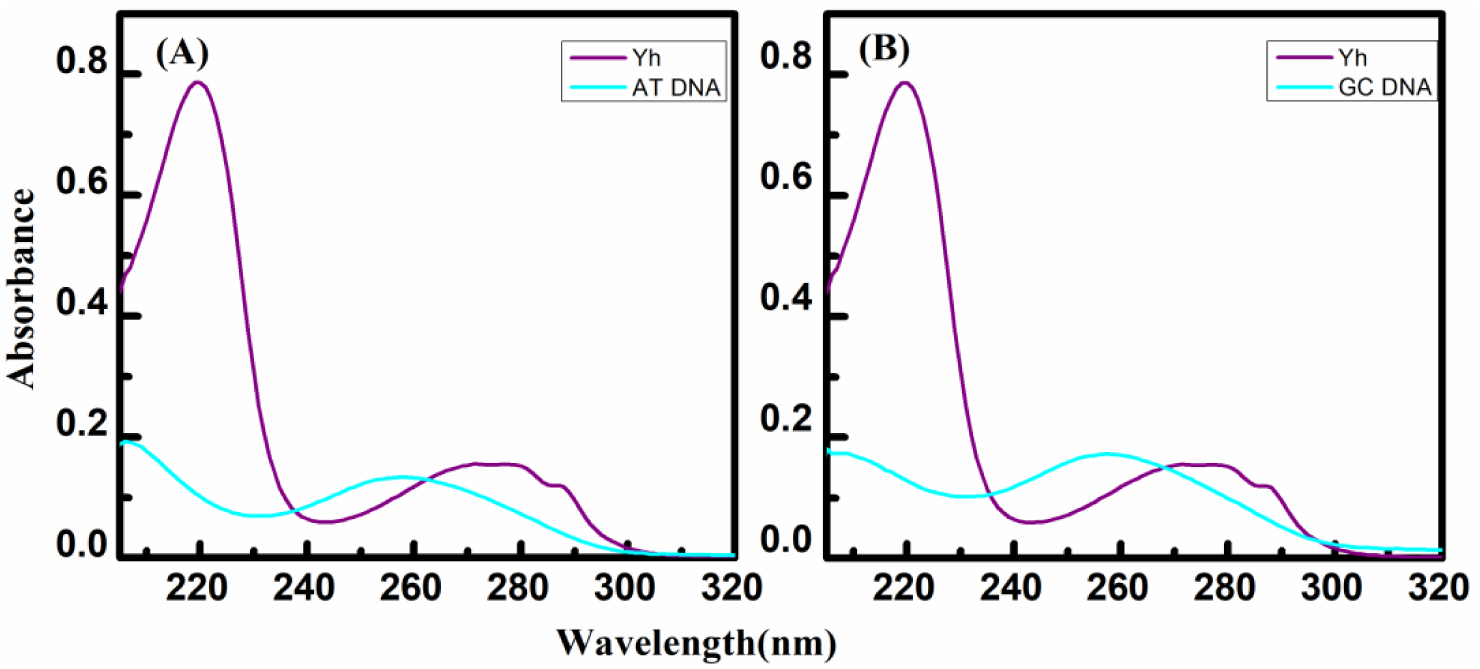
Representative absorption spectral changes of Yh (15 μM) treatment with AT and GC hairpin duplex DNA (15 μM). All experiments were performed in 10 mM sodium cacodylate buffer of pH 7.0 at 298.15 K.

### Steady-state spectroscopy study

For understanding the interaction between ligands (here Yh) and hairpin duplex oligonucleotides, fluorescence spectroscopy has been extensively used [32]. Steady-state fluorescence investigations can be used to generate characteristics like binding affinity, binding sites, dynamics, and conformational alterations. Owing to how weakly DNA fluoresces, Yh was used as a fluorescence probe to ascertain how it interacts with DNA. The spectrum of emission for Yh exhibits a significant intrinsic fluorescence which was in the range from 300 to 440 nm, with a maximum of 352 nm when excited at 250 nm. The emission of fluorescence was quenched by binding to GC and AT hairpin duplex oligonucleotides, which ultimately resulted in the saturation of the site of binding. Figure 3 illustrates the emission patterns for Yh’s interaction with GC and AT hairpin duplex oligonucleotides. Significant emission changes show the ligand has a great affinity for these hairpin duplex oligonucleotide structures, resulting in a strong overlap between the attached ligand and the DNA base pairs. Fluorescence quenching was 74.32% and 62% in GC and AT hairpin duplex oligonucleotide respectively which was not much difference between the two, thus indicating a similar interaction of Yh to both the oligonucleotide structures. With the help of such data, the Scatchard binding equilibrium and further McGhee-von Hippel analysis were used to yield non-linear graphs (inset figure 3)[33]was employed, which revealed non-cooperative binding. Therefore, it is also evident from the fluorescence data that Yh binds to both types of hairpin duplex oligonucleotide (GC and AT) with a little higher affinity. Scatchard binding equilibrium values are employed to reveal the non-cooperative binding; an example of GC and AT hairpin duplex oligonucleotides complexation is displayed as seen in Figure 3 inset. The binding affinity constants were calculated using the fluorescence results according to the Scatchard analysis which utilizes the McGhee-von Hippel analyses’ non-cooperative binding model (*K*) [33],[34]and was found to be 2.1 × 10^5^ M^-1^ and 2.3 × 10^5^ for GC and AT hairpin DNA respectively at 10 mM and 298.15 K temperature. In the case where there is more than one possible binding site in a macromolecule, a double logarithmic equation can be applied to determine the binding constant (*K*) and the number of binding sites. Therefore, the steady-state fluorescence results suggest that Yh binds slightly higher to AT hairpin as compared to GC hairpin duplex oligonucleotides. Its ’*n*’ value, or the most probable number of binding sites, was 2.1 for both the hairpin duplex oligonucleotides.

**Figure 3.**
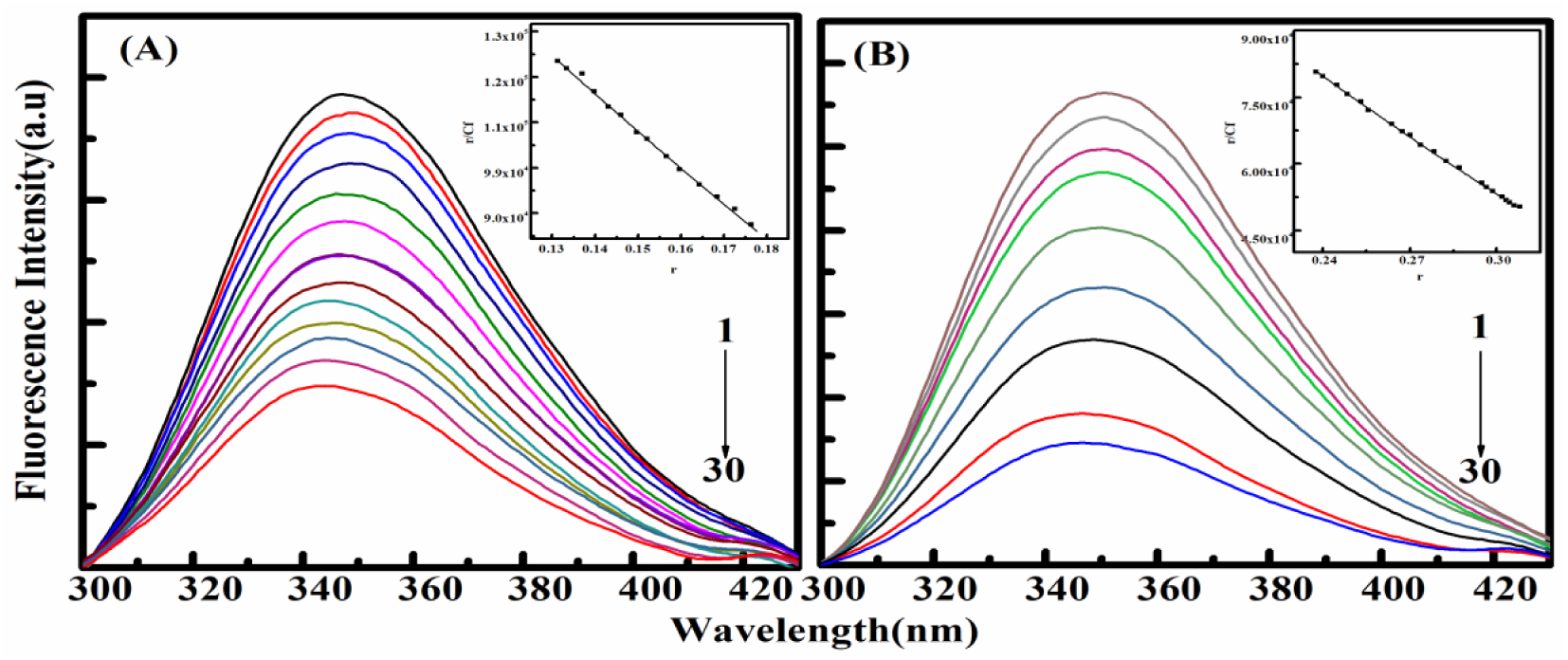
Representative steady-state fluorescence emission spectral changes of Yh (15 μM) with increasing concentration of (A) AT hairpin duplex DNA (0.5 to 14.5 μM) and (B) GC hairpin duplex DNA (0.5 to 14.5 μM) at 298.15 K in 10 mM sodium cacodylate buffer of pH 7.0. Inset: representative Scatchard plot of the binding of Yh with AT and GC hairpin duplex DNA.

### Fluorescence Spectral Study (Temperature-Dependent)

To assess the thermodynamic binding of Yh with GC and AT hairpin duplex oligonucleotide, fluorescence measurement was done at fixed ligand concentrations and escalating concentrations of DNA at 288.15 to 308.15 K. Additionally, it provides a critical understanding of the molecular interactions and the dynamics that promote the development of the complex [35]. The advantage of this method is the fact that all of the interaction’s thermodynamic properties, including stoichiometry and binding affinity constant, can be determined. Figure 3 illustrates represent fluorescence spectra of Yh to GC and AT hairpin duplex oligonucleotide titration at 288.15 to 308.15 K. For temperatures between 288.15 and 308.15 K, Table 1 presents the binding constant of Yh with GC and AT hairpin duplex oligonucleotide. Studies indicated that for interactions between GC and AT hairpin duplex oligonucleotide, the binding affinity constant falls with the increase in temperature. As the temperature increased from 288.15 to 308.15 K, a relatively moderate binding of Yh with GC and AT hairpin duplex oligonucleotide was observed along with a reduction in binding affinity. For Yh with GC hairpin duplex oligonucleotide a fall from 2.4 × 10^5^ M^-1^ to 2.0 × 10^5^ M^-1^ at 308.15 K & for AT hairpin duplex DNA a fall from 2.6 × 10^5^ M^-1^ to 2.1 × 10^5^ M^-1^ at 308.15 K was observed in Table 17. The *K* value obtained from this equation also represents the binding/association constant of the ligand-receptor complex which can interpret whether the binding formed between the DNAs and ligand is strong or weak.

**Table 1.** Binding constant values (*K*) and number of binding sites (*n*) for Yh to AT and GC hairpin duplex DNA interactions at different temperatures. All the experiments were performed in 10 mM sodium cacodylate buffer at pH 7.0.

| **DNAs** | **Temp (K)** | **$K \times 10^5 (\text{M}^{-1})$** | **$n$** |
| --- | --- | --- | --- |
| **AT hairpin duplex** | **288.15** | $2.6 \pm 0.01$ | $1.8 \pm 0.29$ |
| | **298.15** | $2.3 \pm 0.03$ | $2.4 \pm 0.02$ |
| | **308.15** | $2.1 \pm 0.02$ | $1.8 \pm 0.07$ |
| **GC hairpin duplex** | **288.15** | $2.4 \pm 0.08$ | $2.0 \pm 0.03$ |
| | **298.15** | $2.1 \pm 0.25$ | $1.9 \pm 0.02$ |
| | **308.15** | $2.0 \pm 0.06$ | $2.5 \pm 0.31$ |

This is caused by the ligand-DNA complex being destabilized as a result of the increase in temperature. At each of the three temperatures, the value of ’*n*’ (binding site) for GC and AT hairpin duplex oligonucleotide was around 2.0. Even so, the thermodynamic examination of the interactions was rather significant. Regarding Yh to GC and AT hairpin duplex oligonucleotide system, the difference in the Gibbs free energy was relatively minor at all temperatures [36]. Hydrogen bonds, hydrophobic forces, electrostatic interactions, van der Waals forces are the main forces that connect small molecules and biomacromolecules [37]. To determine the basic forces causing a ligand-DNA complexation, one may examine the changes in the free energy (Δ*G°*), enthalpy (Δ*H°*), and entropy (Δ*S°*) for such interactions. Hence, the Gibbs free binding energy (Δ*G°*) derived from the Gibbs-Helmholtz theory and the standard van’t Hoff equation was used to calculate thermodynamic parameters for Yh with GC and AT hairpin duplex oligonucleotide system at 288.15 to 308.15 K:

Δ*H°*,Δ*S°*,and Δ*G°*can be expressed from the following equation (1) and (2).

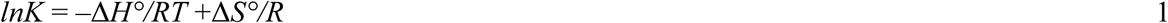

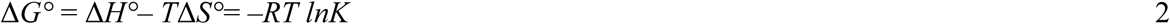

Where R (R = 8.314 J mol^-1^ K^-1^) denotes the universal gas constant and *K* denotes the binding constant at the right temperature. The equation can further be utilized for calculating the free energy change (2). The corresponding values of Δ*H°* and Δ*S°* were calculated from the plot using equation (1) based on *lnK* versus *1/T*, where the slope is equal to –Δ*H°/R* and the point at which it intercepts gives Δ*S°/R* (Figure 4). The resulting values of the parameters Δ*G°*, Δ*H°*, and Δ*S°* were determined using the above equations (1) and (2), and the resulting values are presented in Table 2. According to reports, the main component is found to be electrostatic when Δ*H°*< 0 and Δ*S°*> 0, the significant forces are van der Waals and hydrogen bonding if Δ*H°*< 0 and Δ*S°* < 0, and the essential component is hydrophobic interaction when Δ*H°*> 0 and Δ*S°*> 0 [38],[16].

**Figure 4.**
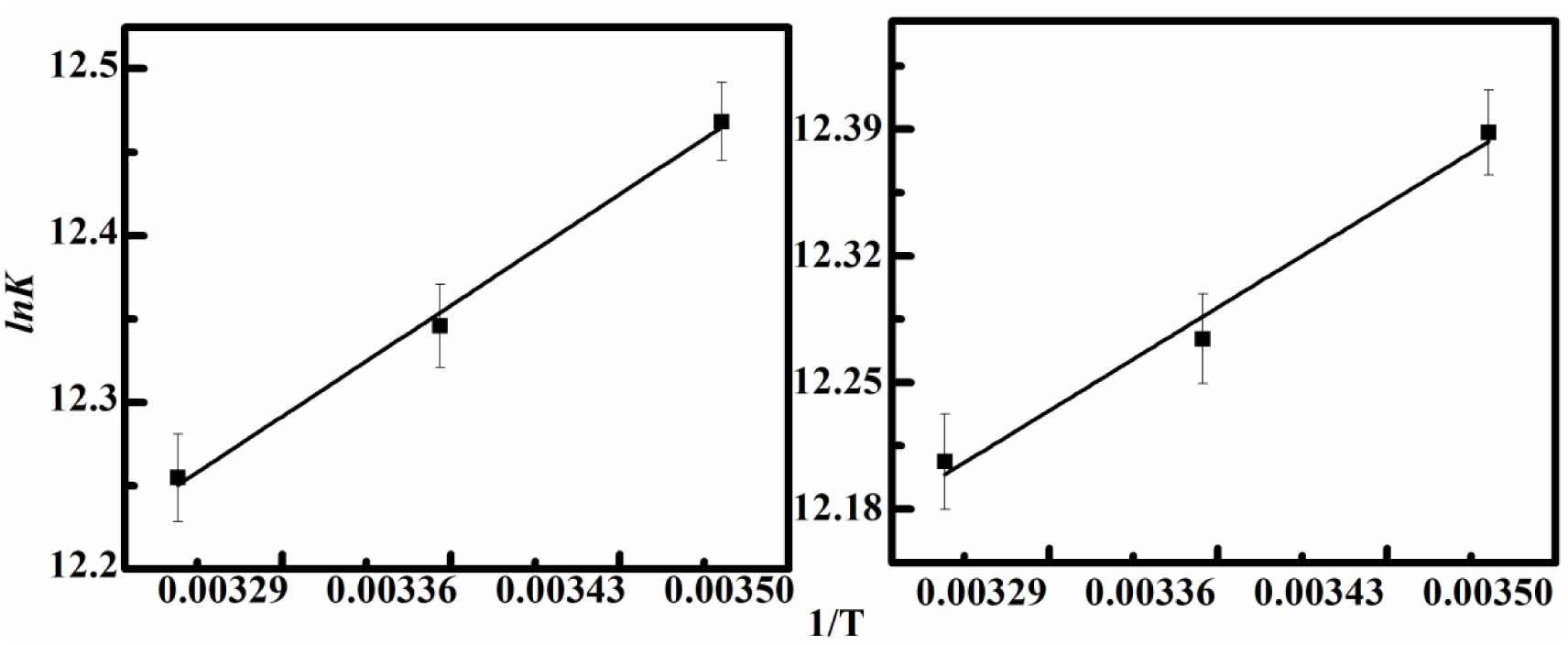
Van’t Hoff Plot of *lnK* versus *1/T* of Yh with (A) AT hairpin duplex DNA and (B) GC hairpin duplex DNA respectively. All experiments were performed in 10 mM sodium cacodylate buffer of pH 7.0 at 298.15 K.

**Table 2.**
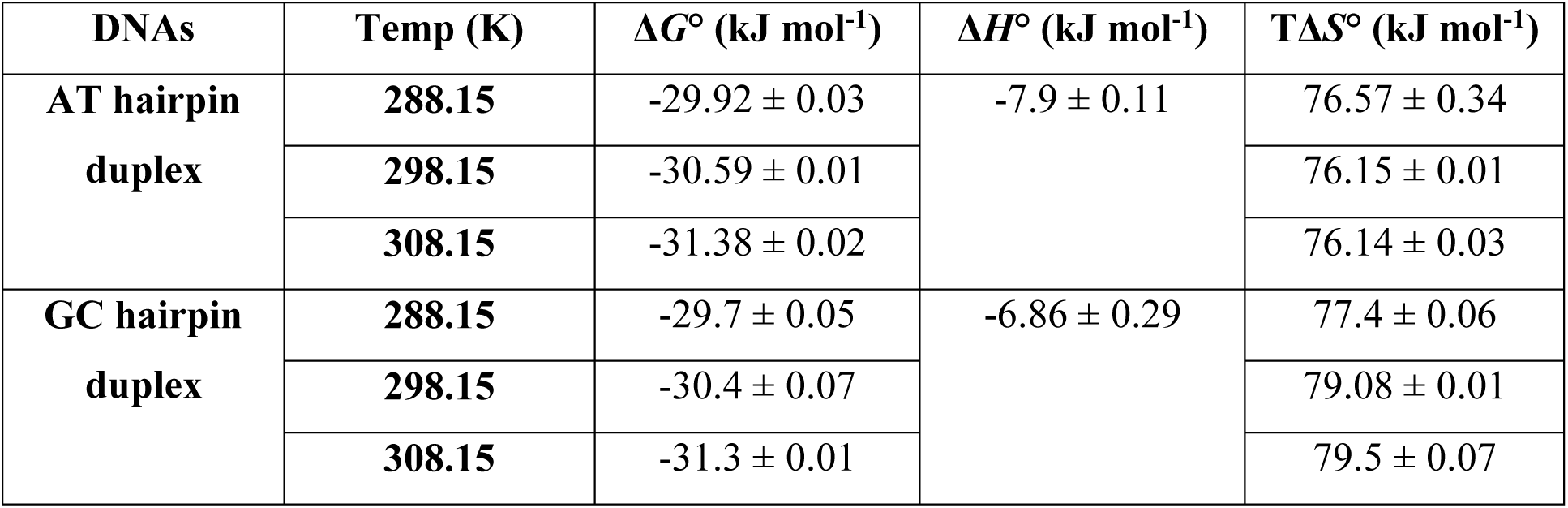
Temperature-dependent thermodynamics parameters for binding of Yh to AT and GC hairpin duplex DNA. All experiments were performed in 10 mM sodium cacodylate buffer of pH 7.0.

| DNAs | Temp (K) | $\Delta G^\circ$ (kJ mol<sup>-1</sup>) | $\Delta H^\circ$ (kJ mol<sup>-1</sup>) | $T\Delta S^\circ$ (kJ mol<sup>-1</sup>) |
| --- | --- | --- | --- | --- |
| **AT hairpin duplex** | **288.15** | $-29.92 \pm 0.03$ | $-7.9 \pm 0.11$ | $76.57 \pm 0.34$ |
| | **298.15** | $-30.59 \pm 0.01$ | | $76.15 \pm 0.01$ |
| | **308.15** | $-31.38 \pm 0.02$ | | $76.14 \pm 0.03$ |
| **GC hairpin duplex** | **288.15** | $-29.7 \pm 0.05$ | $-6.86 \pm 0.29$ | $77.4 \pm 0.06$ |
| | **298.15** | $-30.4 \pm 0.07$ | | $79.08 \pm 0.01$ |
| | **308.15** | $-31.3 \pm 0.01$ | | $79.5 \pm 0.07$ |

As shown in Table 3, the interaction’s thermodynamic analysis between the molecules depicts the binding reaction of both the complex to be in favor of negative enthalpy, Δ*H°*= –6.86kJ mol^-1^& – 7.9 kJ mol^-1^1 (Yh to GC and AT hairpin duplex DNA respectively), and strong positive entropy contribution, Δ*S°*= 79.08 kJ mol^-1^ & 76.15 kJ mol^-1^ for GC and AT hairpin duplex oligonucleotide respectively. As a result, an entropy-driven exothermic binding process is observed [39]. The positive entropy and negative enthalpy are indicators of a DNA-ligand interaction, which often involves a considerable hydrophobic contribution [34]. Gibb’s change in energy for GC and AT hairpin duplex oligonucleotide at 288.15 K was found to be −29.70 kJ mol^-1^ and −29.92 kJ mol^-1^, respectively, while for Yh to GC and AT hairpin duplex oligonucleotide at 308.15 K, it is −31.3 kJ mol^-1^ and −31.38 kJ mol^-1^, respectively. The binding reaction was therefore spontaneous owing to the shift in negative free energy across all temperatures (Δ*G°*< 0).

**Table 3.** Binding constant values (*K*) and number of binding sites (*n*) for Yh to AT and GC hairpin duplex DNA interactions at different salt concentrations. Temperature = 298.15.

| **DNAs** | **$[\text{Na}^+]$ (mM)** | **$K \times 10^5 (\text{M}^{-1})$** | **$n$** |
| --- | --- | --- | --- |
| **AT hairpin duplex** | 10 | $2.3 \pm 0.01$ | $1.8 \pm 0.07$ |
| | 30 | $2.2 \pm 0.04$ | $2.3 \pm 0.01$ |
| | 100 | $1.9 \pm 0.02$ | $2.2 \pm 0.02$ |
| **GC hairpin duplex** | 10 | $2.1 \pm 0.03$ | $1.9 \pm 0.07$ |
| | 30 | $2.0 \pm 0.06$ | $2.0 \pm 0.37$ |
| | 100 | $1.86 \pm 0.08$ | $2.4 \pm 0.09$ |

### Fluorescence Spectral Study (Salt-Dependent)

To deduce the nature of the molecular interactions that control the binding process, salt-dependent fluorescence studies were performed. Along with the van’t Hoff investigation performed in fluorescence at 3 different salt concentrations, it provides details about the chemistry of the interaction study and the influence of the salt concentration on how it binds[40]. Cations exist as counter ions around the DNA double helix, and ligands compete to remove these cations to balance phosphate. The two scenarios are related thermodynamically. It has been determined that as ionic strength increases, the *K* (equilibrium constant) for GC and AT hairpin duplex oligonucleotide decreases. The findings from the salt-dependent fluorimetric investigations are presented in Table 3. As [Na^+^] increased, the *K* values decreased, exhibiting complex formation destabilized at a higher [Na^+^] concentration. Thus, the amount of sodium in the solution directly impacted how strong the interaction was. The *n* values, nonetheless, did not notably change and maintained close to 2, indicating a 2:1 complexation of the two binding species under all salt conditions.

The ligand-DNA binding interactions rely on electrostatic forces. The importance of the binding free energy, as determined by Chaires and his colleagues, is shown by the findings of splitting the binding free energy into 2 constituents: polyelectrolytic and non-polyelectrolytic. We performed a fluorescence study along with van’t Hoff analysis to examine the influence of salt concentrations in the range of [Na^+^] 10 mM to 100 mM on the interaction of Yh to GC and AT hairpin duplex oligonucleotide. The affinity for binding of the interaction decreases as the concentration of [Na^+^] rose from 10 mM to 100 mM [Na^+^]. GC hairpin duplex oligonucleotide, there was a higher drop from 2.1 10^5^M^-1^ at 10 mM to 1.86 10^5^M^-1^ at 100 mM, whereas, for AT hairpin duplex oligonucleotide, it was 2.3 10^5^M^-1^ at 10 mM to 1.9 10^5^M^-1^ at 100 mM (Table 3). The rise in [Na^+^] concentration lowers the electrostatic repulsion between the phosphate groups that are negatively charged with adjacent nucleotides in DNA, which could hinder the ligand’s interaction and result in a decrease in the binding affinity measures shown in (Table 3). As compared to GC hairpin duplex oligonucleotide, the drop in binding capacity for AT was more moderately determined. As a result, the [Na^+^] concentration is thermodynamically associated with the binding of GC and AT hairpin duplex oligonucleotide.

Manning’s counter ions model-based polyelectrolytic theories simply define the process and offer a framework for analyzing the results[41]. The provided equation [42] relates the counter ion release to the slope of the best-fit line in the *lnK* versus *ln[Na^+^]* plot based on the polyelectrolytic theory

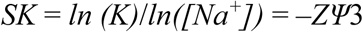

Where Z denotes the apparent charge of the bound ligand per phosphate binding, *Ψ* denotes the proportion of [Na^+^] bound per phosphate group, and *SK* denotes the number of counter ions associated with the binding. The *lnK* versus *ln[Na^+^]* slope (Figure 5) gave values of 0.13. The results of this study show identical electrostatic interactions between Yh and GC and AT hairpin duplex oligonucleotide and represent the counter ions released per phosphate of DNA during binding. Similar support regarding this is offered by partitioning the observed binding Gibbs free energy into contributions from polyelectrolytic (Δ*G°_pe_*) and non-polyelectrolytic (Δ*G°_t_*) systems. The relationship described below can be used to determine the polyeletrolytic contribution to the observed variation in Gibbs free energy

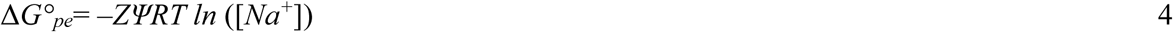

**Figure 5.**
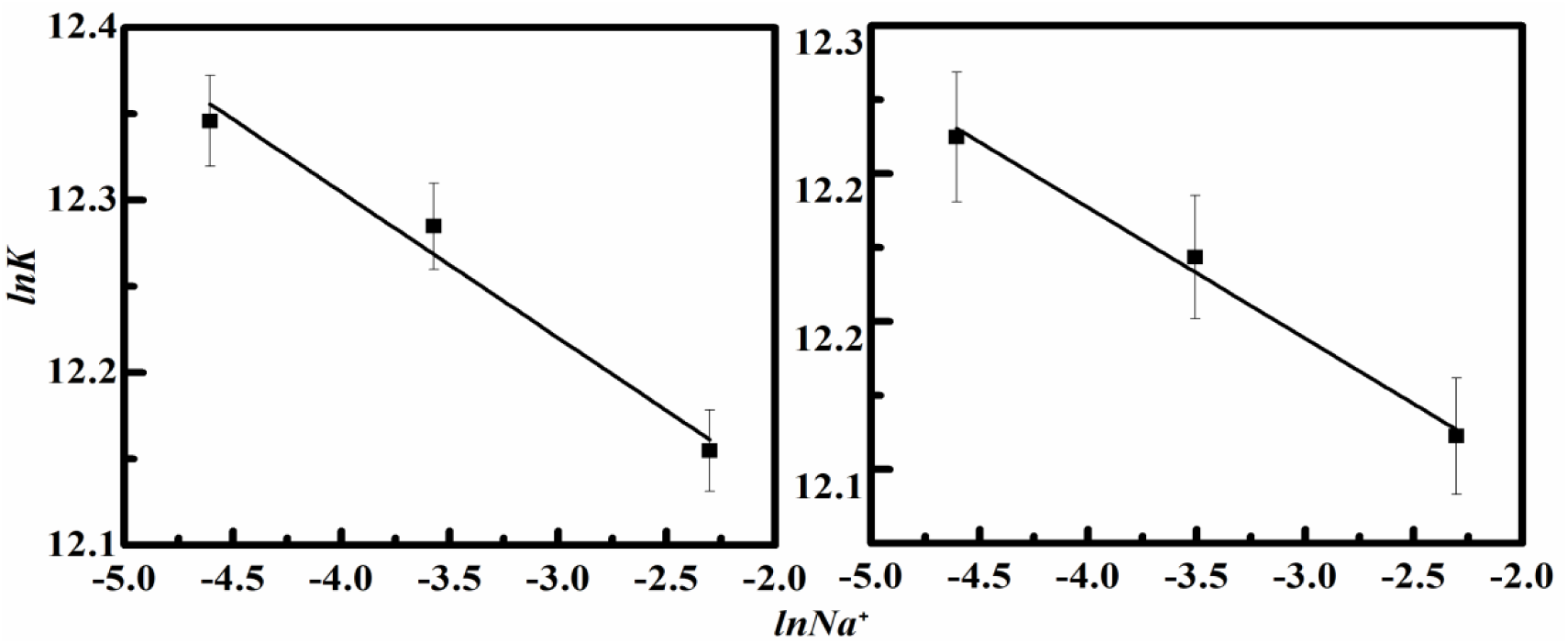
Van’t Hoff Plot of *lnK* versus *ln[Na+]* of Yh with (A) AT hairpin duplex DNA and (B) GC hairpin duplex DNA respectively. All experiments were performed at 298.15 K.

The non-polyelectrolytic Δ*G°_t_* contribution is determined by the difference between Δ*G°* and Δ*G°_pe_* at a particular concentration of [Na^+^]. Here, ZΨ represents the slope of the van’t Hoff plot (see Figure 6). At 10 mM [Na^+^], the contribution from Δ*G°_pe_* is calculated to be −0.57 kcal mol^-1^ & −1.03 kcal mol^-1^ which was about 7.85% & 14.48% for GC and AT hairpin duplex DNA respectively, from the total change in Gibbs free energy. Salt concentration at 100 mM [Na^+^], the values of Δ*G°_pe_* were determined to be −0.29 kcal mol^-1^ & −0.51 kcal mol^-1^ which was about 4.03% & 7.36% for GC and AT hairpin duplex DNA respectively.

**Figure 6.**
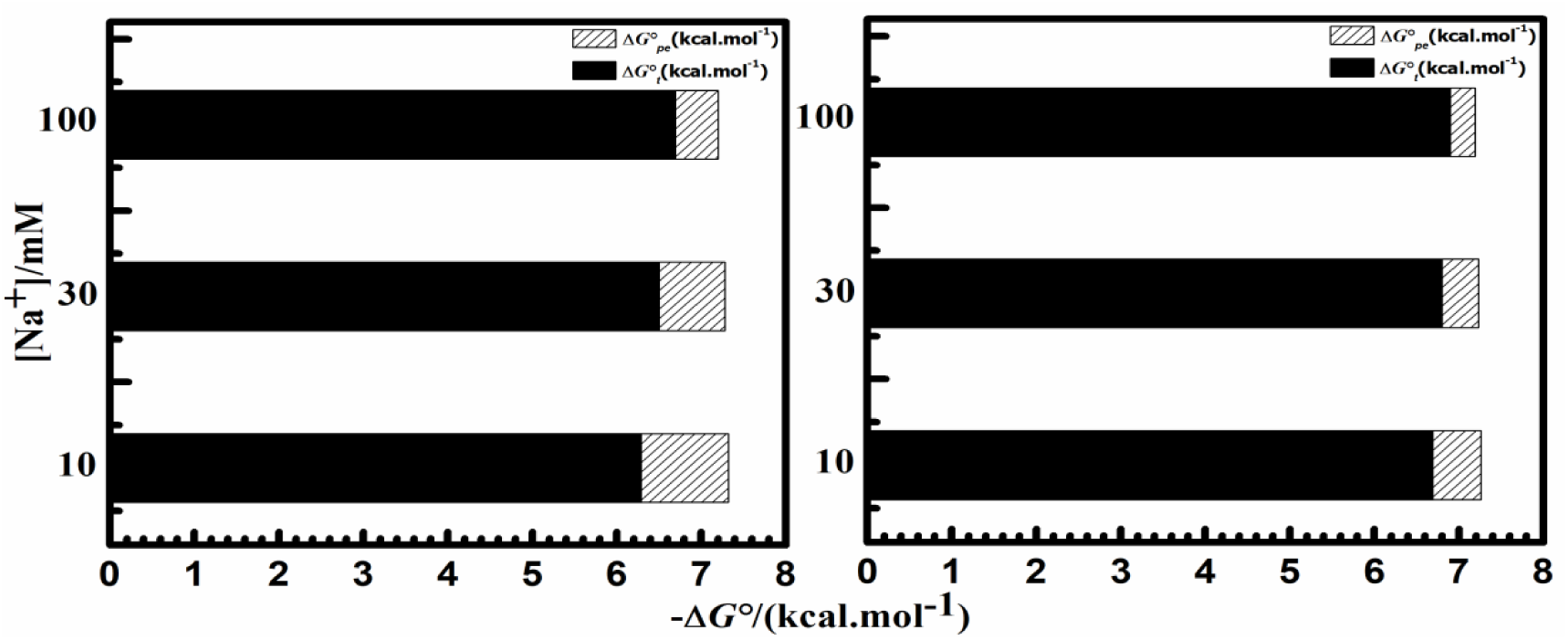
Partitioned polyelectrolytic (*ΔG°_pe_*) (shaded) and non-polyelectrolytic (*ΔG°_t_*) (black) contribution to the standard molar Gibbs energy at (10, 30, and 100) mM [Na^+^] concentrations (A) AT hairpin duplex DNA and (B) GC hairpin duplex DNA respectively. All experiments were performed at 298.15 K.

Figure 6 and Table 4 provide an example of the partitioned change in Gibbs free energy. It may be observed that in each of the cases when there is an increase in [Na^+^] concentration, the Δ*G°_t_* contribution remained constant while the contribution of Δ*G°_pe_* decreased. Non-polyelectrolytic (Δ*G°_t_*) is primarily responsible for stabilizing the formation of the complex between Yh binding to GC and AT hairpin duplex oligonucleotides. Thus, the results presented here confirm the contribution of non-polyelectrolytic forces to the binding interaction. The overall binding of Yh to GC and AT hairpin duplex oligonucleotide has a non-polyelctrolytic contribution a lot higher than the polyelectrolytic contribution.

**Table 4.** Salt-dependent thermodynamics parameters for binding of Yh to AT and GC hairpin duplex DNA. All experiments were performed at a temperature = 298.15.

| DNAs | $[Na^+]$ (mM) | $\Delta G^\circ$ kcal/mol | $\Delta G^\circ_t$ kcal/mol | $\Delta G^\circ_{pe}$ kcal/mol | $Z\Psi$ |
| --- | --- | --- | --- | --- | --- |
| AT hairpin duplex | 10 | $-7.11 \pm 0.01$ | $-6.29 \pm 0.27$ | $-1.03 \pm 0.03$ | $0.09 \pm 0.30$ |
| | 30 | $-6.96 \pm 0.02$ | $-6.5 \pm 0.01$ | $-0.78 \pm 0.06$ | |
| | 100 | $-6.93 \pm 0.08$ | $-6.69 \pm 0.03$ | $-0.51 \pm 0.25$ | |
| GC hairpin duplex | 10 | $-7.26 \pm 0.25$ | $-6.69 \pm 0.27$ | $-0.57 \pm 0.08$ | $0.05 \pm 0.03$ |
| | 30 | $-7.23 \pm 0.01$ | $-6.8 \pm 0.03$ | $-0.43 \pm 0.31$ | |
| | 100 | $-7.19 \pm 0.06$ | $-6.9 \pm 0.01$ | $-0.29 \pm 0.29$ | |

### Potassium iodide (KI) quenching assay

Potassium iodide (KI) which is a quenching compound was used to decode how Yh ligand interacts and binds with GC and AT hairpin duplex oligonucleotide. DNA has a framework of negatively charged phosphate which is expected to repel or oppose a quencher which is extremely negatively charged, intercalated small molecules in DNA’s double helix act as barriers from an ionic quencher’s effects. However, anionic quenchers readily quench the molecules that bind the DNA via groove-binding mode and it remains in contact with the nearby surroundings [43],[44]. Therefore, on treatment with KI, the variation in K_sv_ values between groove-binding molecules and intercalating molecules is larger [45]. A total of three experiment sets were carried out to study the fluorescence spectra of Yh alone, Yh‒AT, and Yh‒GC hairpin duplex oligonucleotide complexes after the titration with Potassium iodide (KI). Using the stated formula that follows the equation, the Stern-Volmer (K_sv_) values were determined from the Stern-Volmer plot (Figure 7):

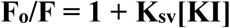

**Figure 7.**
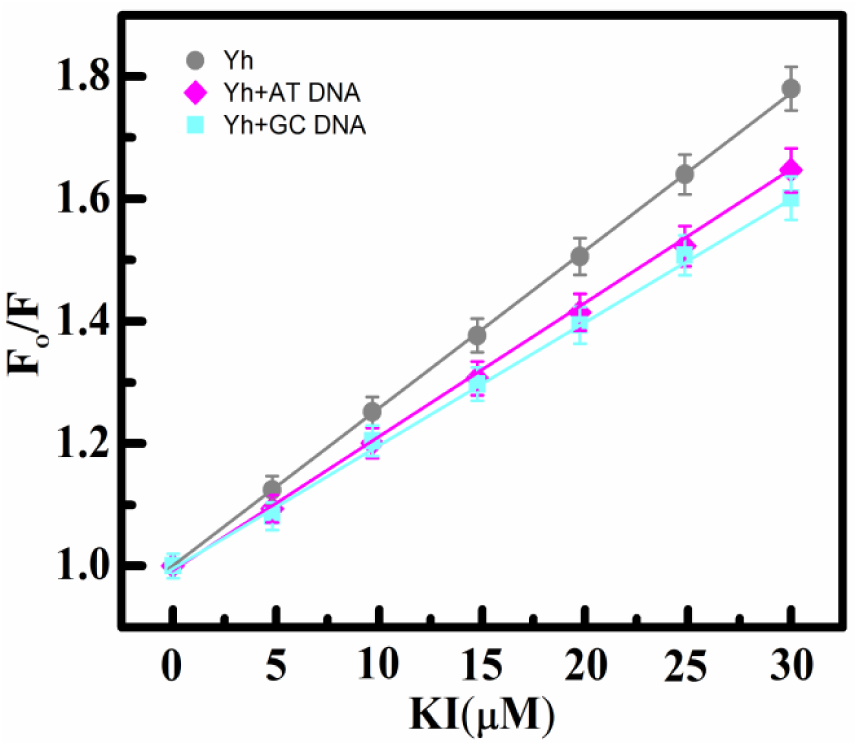
The Stern-Volmer plot of fluorescence intensity of Yh (20 μM) titrated with KI (0-30 μM) in the absence and presence of AT and GC hairpin duplex DNA (20 μM). Temperature = 298.15 K. All experiments were performed in 10 mM sodium cacodylate buffer of pH 7.0.

Where F and F_o_ stand for the fluorescence intensities in the presence and absence of the anionic quencher here potassium iodide (KI), respectively. The K_sv_ values of Yh alone, Yh-AT, and Yh-GC hairpin duplex oligonucleotide were determined to be 1.3 × 10^3^ M^-1^, 1.02 × 10^3^ M^-1^, and 1.05 × 10^3^ M^-1^ respectively. K_sv_ (Stern-Volmer constant) obtained from Stern-Volmer equation helps determine the type of quenching mechanism involved between the fluorophore and the quencher. It gives information about the quenching efficiency and the rate of quenching. A higher Stern-Volmer constant (K_sv_) implies a more vital quenching interaction between the fluorophore and quencher. For binding of Yh to AT and GC DNA, the K_sv_ value of the free Yh was higher than that of the bound Yh to AT or GC indicating groove binding. The result displayed almost the same iodide quenching effect without much difference in the fluorescence of Yh before and after the interaction with GC and AT hairpin duplex oligonucleotide. Thus, supporting the groove binding mode of Yh with both the DNA [45],[46].

### Urea-induced denaturation study

This type of study is also carried out to confirm the way small molecules bind to hairpin duplex oligonucleotides. Small intercalated compounds are released from DNA as a result of compounds that act as denaturants (urea) destabilizing the native dsDNA helix structure, which changes the small molecule’s fluorescence intensity [47]. The fluorescence intensity of groove-binding ligands is practically unchanged with the existence of urea.

Yh-AT and Yh-GC hairpin duplex oligonucleotide interactions were titrated with urea, and their fluorescence spectra were obtained (Figures 8A & B). Upon titrating with urea, it became evident that there is no noticeable difference in the intensity of fluorescence, which indicates that Yh binds to GC and AT hairpin duplex oligonucleotide in a groove region instead of intercalation[48]. Figures 8C & D illustrate the association linking the fluorescence intensities of Yh‒AT and Yh‒ GC hairpin duplex oligonucleotide interacting complex in the presence and absence of denaturant (here urea) (F_o_/F). Therefore, our work supports Yh’s groove binding mode with GC and AT hairpin duplex oligonucleotide additionally.

**Figure 8.**
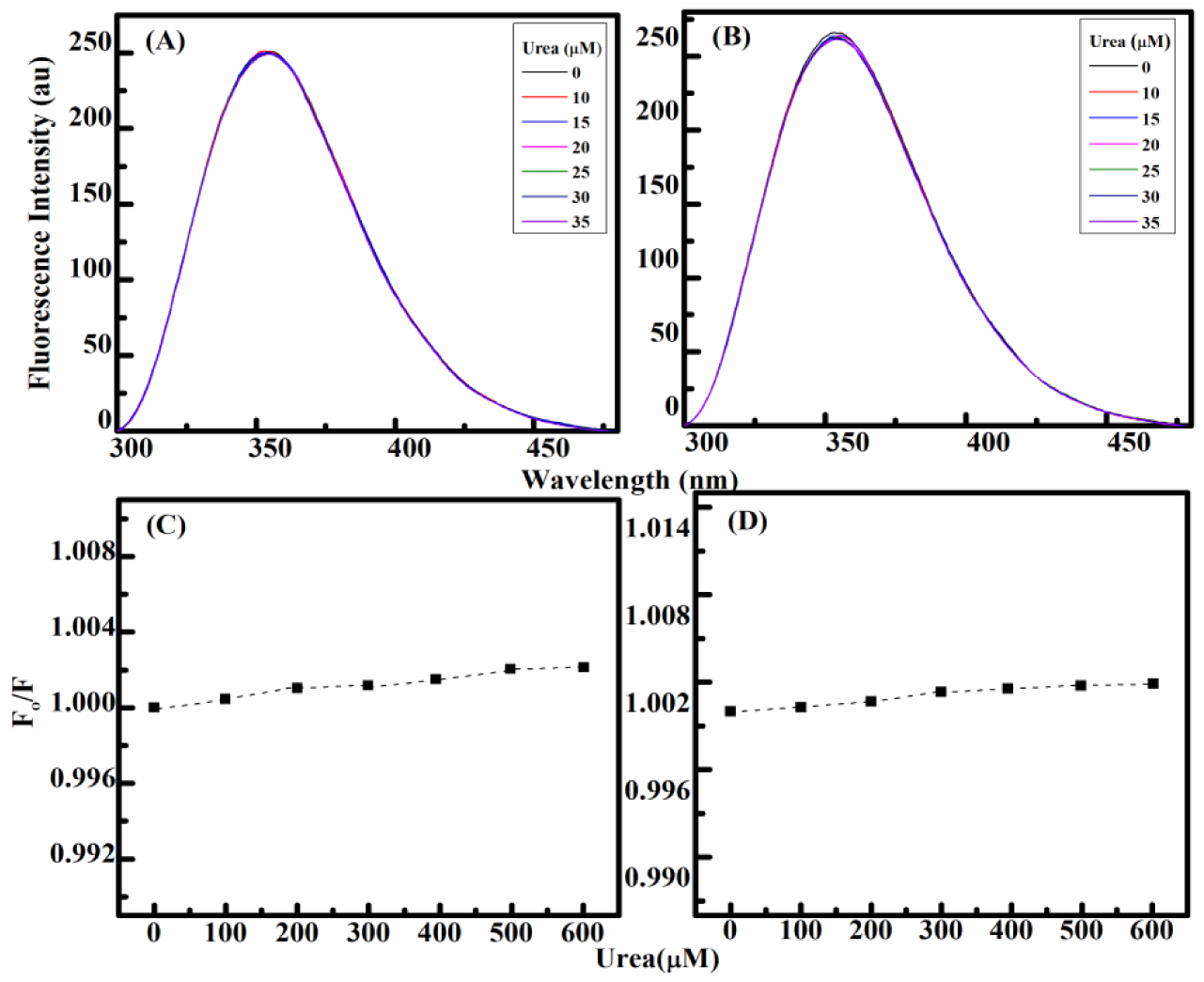
Fluorescence emission spectra of Yh with (A) AT DNA and (B) GC DNA titrated with urea (0-35 μM) respectively. The plot of F_o_/F versus concentration of urea (C) AT DNA and (D) GC DNA. Temperature = 298.15 K. All experiments were performed in 10 mM sodium cacodylate buffer of pH 7.0.

### Competitive drug displacement assay

Each of these fluorescence-based assays was performed to prove how ligands bind to DNA. Fluorescent dyes with confirmed DNA binding modes are utilized [49],[50]. For example, ethidium bromide (Etbr) binds to dsDNA by mode of intercalation whereas rhodamine-B binds through a groove-binding mode of interaction [51],[52],[53].

Rhodamine-B is one of the common groove binder indicators and ethidium bromide is a common intercalative binder indicator. Rhodamine-B and ethidium bromide (Etbr) were used as markers in the dye displacement binding assay to support the mode of binding mechanism of Yh with GC and AT hairpin duplex oligonucleotide. Dye-DNA complexation is formed to interact with Yh, where the binding mechanism needs to be determined, with a shift in the dye-DNA complexation intensity of fluorescence monitored. There is speculation that the DNA helix will interact with the Yh, which competes and eliminates the dye attached to hairpin oligonucleotides, like the displaced dye [50].

The dye-GC and AT hairpin duplex oligonucleotide complexes were titrated by higher concentrations of Yh in multiple sets of experiments, while the resulting fluorescence spectra were obtained. When GC and AT hairpin duplex oligonucleotide complexes with Rhodamine-B were titrated with Yh, a decrease in the intensity of fluorescence was noticed (Figure 9B & D), but no such impact was seen when ethidium bromide was used (Figure 9A & C). For both the hairpin duplex oligonucleotides, Rhodamine-B variations in fluorescence intensity were significantly greater compared to that of ethidium bromide (Figure 9) [54],[46]. These observations confirm that Yh binds to GC and AT hairpin duplex oligonucleotide via a groove binding interaction which was previously mentioned in the experimental findings [55],[56].

**Figure 9.**
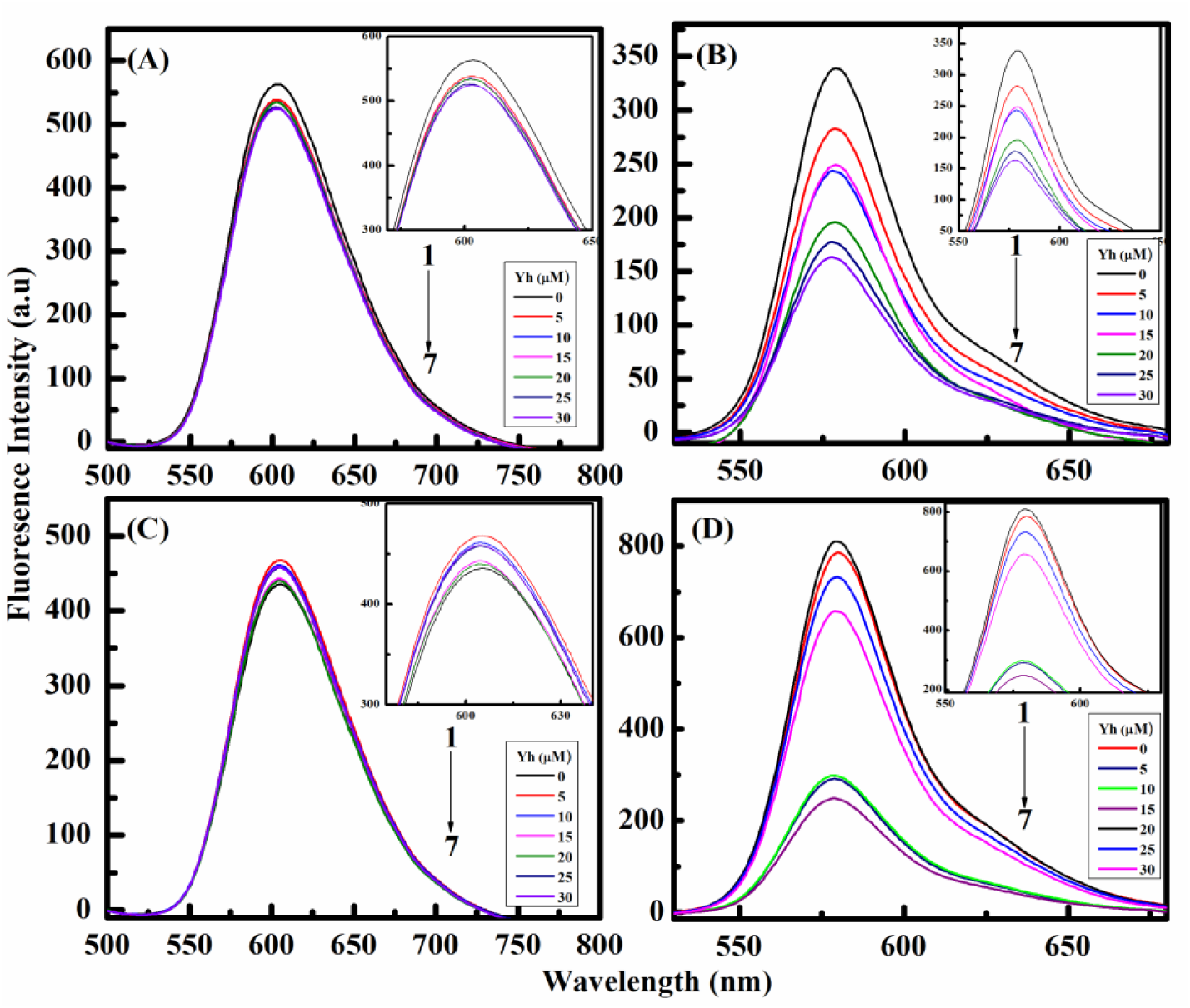
Fluorescence emission spectra of Ethidium bromide with (A) AT DNA and (C) GC DNA titrated with Yh (0-30 μM) and Rhodamine-B with (B) AT hairpin duplex DNA and (D) GC hairpin duplex DNA titrated with Yh (0-30 μM) respectively. Temperature = 298.15 K. All experiments were performed in 10 mM sodium cacodylate buffer of pH 7.0.

### CD Spectral study

Circular dichroism spectroscopy is an effective tool for evaluating the native structural feature of DNA when small molecules interact with different types of DNA, such as A, B, Z, etc. CD signals are a powerful and quite sensitive tool for the determination of binding interaction patterns [57][58]. A very prominent positive signal at 275 nm caused by DNA base-pair stacking and a negative signal at 245 nm caused by its helical configuration (indicating DNA as a right-handed B configuration) were visible in the CD spectra [57],[59]. It needs to be highlighted that the B form of DNA is considered a primary physiological form of dsDNA[60][61]. These specified bands have been suggested as highly responsive to ligands interacting with DNA. CD spectrum for the solution of GC and AT hairpin duplex DNA (10 μM) in the presence and absence of Yh (0 μM-60 μM) are shown in Figure 10. The structure of native DNA may perturb on interaction with small molecules like Yh. The level and form of interaction determine the rate of the shifts [62],[63]. The naturally present right-handed B-form of DNA is stabilized by the ligand molecules when it intercalates to DNA, which results in the multiplication and shifting of both bands. Similarly, electrostatic binding and basic groove binding also cause minimal to no band shifts [64]. The minor disturbance caused by the positive peak and the negative band, suggests that the addition of Yh is not significantly affecting the helicity of GC and AT hairpin duplex oligonucleotides [65]. This shows that Yh to GC and AT hairpin duplex oligonucleotide binding causes some minor perturbation in the DNA molecule, such as the transition from B to C-like structure [66]. These minor modifications indicate Yh’s non-intercalative binding which also supports the groove binding nature [67]. The finding presents and supports the earlier evidence analyzed by using various experiments.

**Figure 10.**
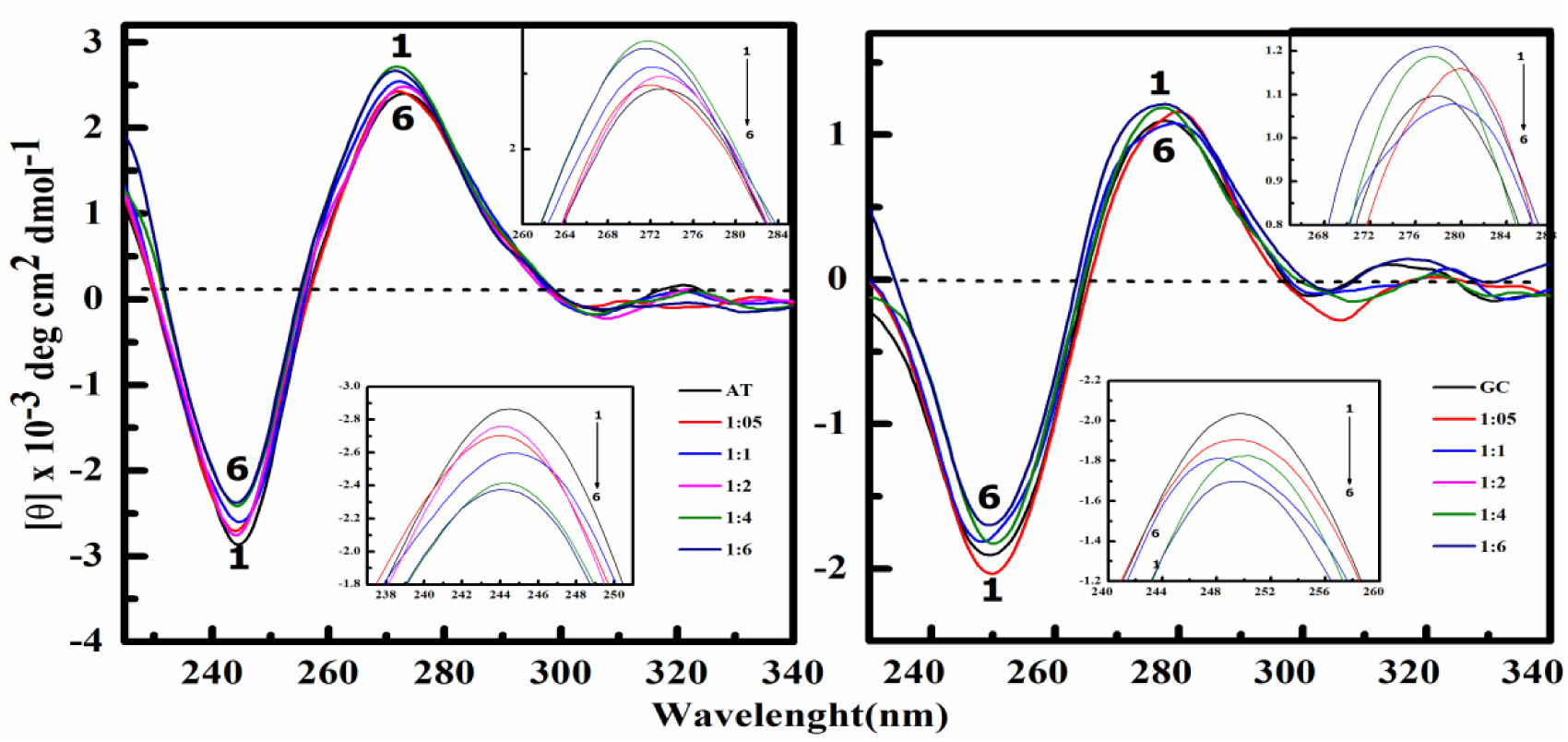
CD spectra for the interaction of (A) AT hairpin duplex DNA (10μM) and (B) GC hairpin duplex DNA (10μM) treated with 5, 10, 20, 40, and 60 μM (curves 1-6) of Yhat 298.15 K in 10 mM sodium cacodylate buffer of pH 7.0.

### Molecular modeling studies

An insight into the ligand’s interactions with hairpin oligonucleotide is revealed via *in silico* molecular modeling analysis. AutoDock 1.5.6 program was applied to dock the conformer with the lowest energy of Yh with GCandAT hairpin duplex oligonucleotide (PDB-ID: 5M68, 1D16) to explore the binding associations [68],[69]. In all cases, docking was performed to clarify the forces involved and better comprehend how Yh binds to GC and AT hairpin duplex oligonucleotide. Figures 11A & B show that the Yh molecule visibly interacts with both the oligonucleotide hairpin duplex. When Yh binds to GC and AT hairpin duplex oligonucleotide (5M68, 1D16), the electrostatic binding energy is considerably smaller than the total of van der Waal’s energy, hydrogen bonding, and desolvation-free energy. In every case, hydrogen bonding (H-bonding) was present. At a distance of 2.106 Å, the carboxylate component of Yh constituted a hydrogen-bond with the sugar-phosphate backbone of AT hairpin duplex oligonucleotide, however at a distance of 1.77 Å; the carboxylate portion of Yh formed a hydrogen bond with the keto group of thymine for GC hairpin duplex oligonucleotide (Figure 12).

**Figure 11.**
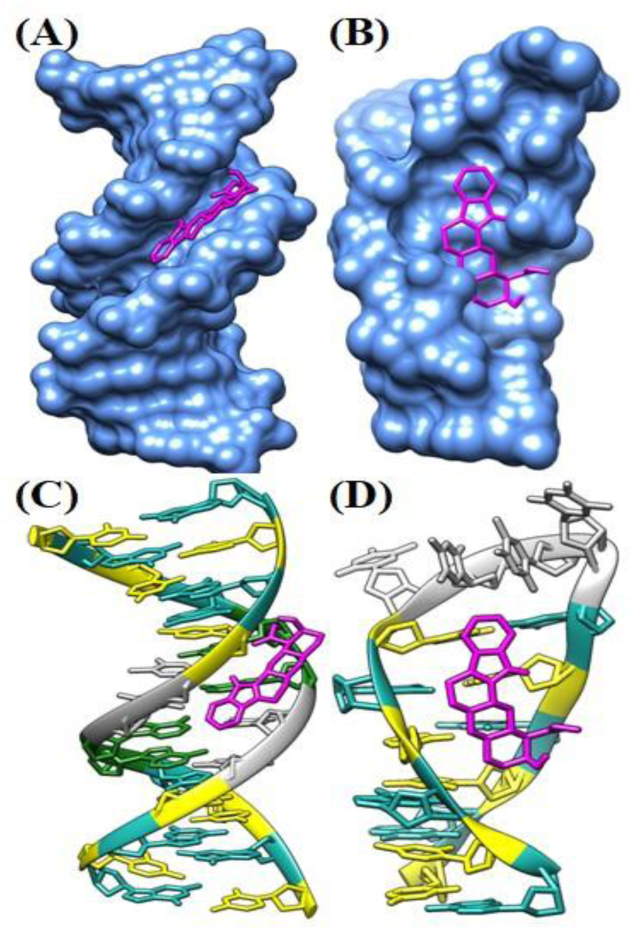
(A)&(B) Docked pose of Yh with AT and GC hairpin duplex DNA (PDB ID: 5M68, 1D16), (C)&(D) Hydrophobic surface of AT and GC hairpin duplex DNA (5M68, 1D16).

**Figure 12.**
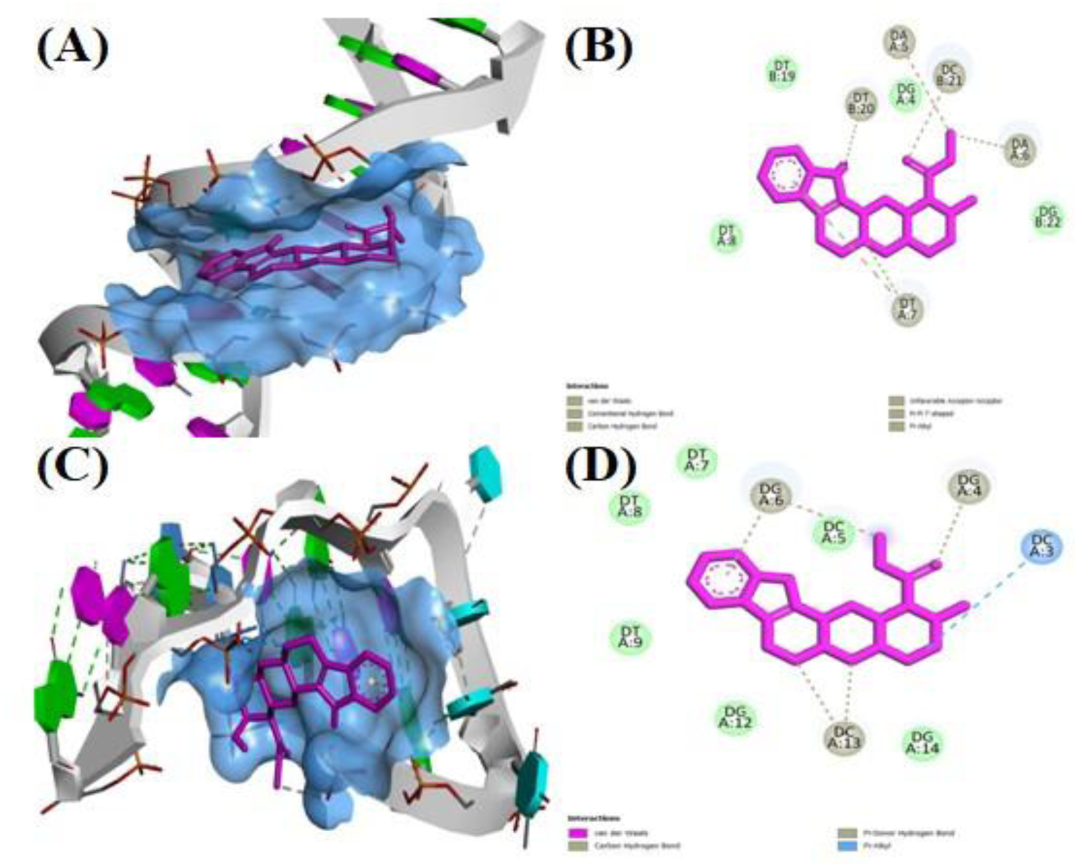
(A)&(B) Hydrophobic forces covering the surface of AT and GC hairpin duplex DNA (5M68, 1D16) with Yh, (C) & (D) 2D diagram indicating the DNA base pairs of AT and GC hairpin duplex DNA (5M68, 1D16) with Yh and interacting forces in the complex formation.

The binding energy which was predicted due to the interaction of Yh with GC and AT hairpin duplex oligonucleotide is −9.35 kcal mol^-1^ and −9.89 kcal mol^-1^ respectively. The binding energy resulting from molecular docking analysis complements the experimental data. Inhibition constant Yh was 139.71μM & 56.06μM for GC and AT hairpin duplex DNA respectively. Thus, groove binding emerged as a means of binding, which also supports the experimental results.

## Conclusion

Yohimbine, an indole alkaloid drug, was investigated for its interactions with GC and AT hairpin duplex oligonucleotide. Yh with GC and AT hairpin duplex oligonucleotide is quantitatively explained through the usage of various biophysical tools, which include both spectroscopy and computational approaches. By UV-Vis and Steady-state spectroscopy, the binding process resulted and was directed toward the occurrence of a specific ligand-DNA complex (that includes 2:1 stoichiometry) at the ground state. The binding affinities determined by the methods were 10^5^ M^-1^ for both cases. Temperature-dependent fluorescence thermodynamics features suggest the binding process is an exothermic reaction driven by entropy with a spontaneous Gibb’s free energy value; the entropy and enthalpy data indicated hydrophobic and electrostatic interactions. The result also determined that non-polyelectrolytic contribution was dominant. Potassium iodide, urea denaturation, and dye displacement assay showed the groove binding mode for both the hairpin duplex DNA. The CD spectra revealed that the ligand Yh caused a slight conformational alteration in both hairpin duplexes, which may be a groove binder. However, the obtained stoichiometry and binding constant complemented the spectroscopic data. The comprehensive interaction pattern is illustrated by molecular docking simulation studies, highlighting the role of H-bonding and hydrophobic interactions in determining the binding energy. The present research provides a quantitative analysis of a plant-derived bioactive interaction with hairpin duplex oligonucleotide by detailed thermodynamic and structure-function relationship analysis. Such bindings could change the DNA’s structural stability and thus disrupt its biological function.

## Author’s Contributions

Concept, design, and overall guidance of the research: J.B; experiment, data analyses, writing/draft preparation: S.L. partial experimental analysis: S.Y. partial writing/draft preparation: V.R. All the authors have read and approved the final version of the manuscript.

## Supporting information

ESI

## Acknowledgment

Financial assistances from DBT, Govt. of India (Twinning research scheme; Sanction Order No. BT/PR25026/NER/95/963/2017)and DRDO, Govt. of India (funding through North East Science & Technology Centre, Mizoram University; Project no. DFTM/07/3603/NESTC/EWM/P-04)) are thankfully acknowledged. The authors would like to thank Dr. Souvik Maiti, Director and Senior Principal Scientist, CSIR – Institute of Genomics and Integrative Biology, New Delhi, for his throughout support.

## Declaration of competing interest

The authors declare that they have no known competing financial interests or personal relationships that could have appeared to influence the work reported in this paper.

## Notes

### Competing Interest Statement

The authors have declared no competing interest.

