## Supplementary material for "Study on the interaction of Yohimbine with duplex oligonucleotide using spectroscopic and computational tools": ESI

### Electronic Supplementary Information (ESI)

### Methods

#### UV-Vis Absorption Spectroscopy

An Agilent, Cary 100 series UV-Vis spectrophotometer in standard quartz cuvettes of 3.5 ml with a 10 mm optical path length was used to analyze the absorption spectral studies at (298.15 ± 0.5) K. For the ligand–DNA study, before recording the absorbance values, the solution was properly mixed and allowed to change state after each fraction of the drug was added to the DNA solution. At a minimum, three continuous measurements were taken for each sample and averaged out.

#### Steady-state spectroscopy study

Steady state fluorescence spectra were measured at (298.15 ± 0.5) K on an Agilent Cary Eclipse spectrofluorimeter. The fluorescence measurements employed 10 mm optical path length standard quartz cuvettes. For measurements of the intrinsic fluorescence spectral change of Yh due to the complexation with increasing AT and GC hairpin duplex oligonucleotide concentration, the ligand samples (dissolved in buffer) were excited at 250 nm. The excitation maxima of Yh and the emission spectra scanned in the range of 300 to 440 nm. At least three consecutive measurements were taken for each sample and averaged out.

#### Analysis of Binding Data

Titration data were used to contrive Scatchard plots, *i.e.* by plotting $r/C_f$ versus $r$ for investigation; in which $r$ is the number of drug molecules bound per mole of DNA and $C_f$ is the free Yh concentration. In the case of normal titrations, the concentrations of free Yh ($C_f$) and bound Yh ($C_b$) are calculated using $C_f = C(1 - \propto)$ and $C$b ) $C_T - C_f$, respectively, where $C_T$ is the total Yh concentration (15 $\mu$M). The bound fraction of Yh ($\propto$) was calculated using the equation, $\propto = (A_f - A)/(A_f - A_b)$, where $A_f$ and $A_b$ are the absorbance of the free and fully bound Yh at the absorption maxima of Yh *i.e* at 352 nm, respectively, and $A$ is the absorbance of Yh at 352 nm at any given point during the titration. As the binding for the interaction of Yh with different AT and GC hairpin duplex oligonucleotide obtained here was nonlinear, to calculate the binding parameters with higher accuracy, nonlinear fitting models were used as described below.

Data were fitted using the nonlinear Scatchard equation:

$$r/C_f = K \times (1- n \times x) \times ((1- n\times x)/(1- (n-1) \times x))^{(n-1)}$$

where $r$ is the number of alkaloid molecules bound per mole of DNA base pair and $C_f$ is the molar concentration of the unbound alkaloids, $n$ is the no of alkaloids bound per DNA base pair and $K$ represents the affinity of the ligand's for the binding site. Further the binding data were plotted as $r$ vs $C_f$ and analyzed by nonlinear curve fitting; Origin 8.5 was used for all the fitting analyses.

**Potassium iodide (KI) quenching experiments**

The quenching effect of KI was studied by adding stoichiometric small aliquots of potassium iodie (KI) stock solution to the three sets of the experiment. In one set of experiments, 20 μM of Yh was titrated with KI (0-30 μM). In another set, the Yh-AT and GC hairpin duplex oligonucleotide complex (2:1 ratio) was titrated with KI (0-30 μM). The fluorescence intensity data were recorded, and then the quenching constants were calculated to form Stern-Volmer plots.

**Urea-induced denaturation study**

This assay was done in three experimental setups. In the first experiment, 15 μM of Yh was titrated with urea (0-35 μM). In another one, the Yh to AT and GC hairpin duplex oligonucleotide complex (2:1 ratio) was titrated with urea (0-35 μM) to record the emission spectra.

**Competitive drug displacement assay**

The competitive interaction between rhodamine B and Yh with AT and GC hairpin duplex oligonucleotide was carried out as follows: fixed amounts of rhodamine B (10 μM) and AT and GC hairpin duplex oligonucleotide (10 μM) were titrated by successive additions of Yh (0-30 μM). The emission was observed between 400-700 nm after exciting at 350 nm. The competitive interaction between ethidium bromide and Yh with both the hairpin duplex oligonucleotide was carried out as follows: fixed amounts of ethidium bromide (10 μM) and AT and GC hairpin duplex oligonucleotide (10 μM) were titrated by successive additions of Yh (0-30 μM). The emission spectra were observed between 500-800 exciting at 475 nm.

**CD spectral study**

CD experiments were performed with the JASCO 1500 spectropolarimeter (JASCO, Japan) under a constant flow of nitrogen. All measurements were done at sodium cacodylate buffer (10mM), 298.15 K with a 1 mm quartz cuvette, and covering a spectral range of 200-400 nm using a scan rate of 200 nm/min, 1 s response time, and 1 nm bandwidth. The spectra were measured with a fixed duplex DNA concentration (10 μM) and a variation of drug concentration (5-60 μM). Each spectrum was obtained as an average of three measurements and was baseline corrected.

**Molecular modeling studies**

AutoDock version 1.5.6 was used to do the molecular docking. Yh interaction with AT and GC hairpin duplex oligonucleotide is investigated in this docking. According to the literature data, it could be assumed that the similar GC content of AT (PDB ID: 5M68) and GC (PDB ID: 1D16) hairpin duplex oligonucleotide, and crystal structure were provided by the Protein Data Bank. The complete water molecule was deleted for both the DNA molecule construction, and polar hydrogen and Gasteiger charges were inserted into the macromolecule file. To contain the complete DNA molecule, the grid spacing was set at 0.375 Å with the dimension of 98 Å × 126 Å × 68 Å (X-Y-Z) for both the hairpin duplex oligonucleotide. The Lamarckian Genetic Algorithm was employed as a docking parameters algorithm in the docking operation, and some other parameters were set to the standard values provided by AutoDock 1.5.6. Based on the docked configuration with the lowest amount of energy, as calculated by the AutoDock scoring tool, the active binding was selected for each docking condition. PyMOL (The PyMOL Molecular Graphics System, Version 2.3.4, Schrödinger, LLC), UCSF Chimera 1.15 molecular graphics tool, and Discovery studio were used to visualize the docked position.
